# Increased Dynamics of α-Synuclein Fibrils by β-Synuclein Leads to Reduced Seeding and Cytotoxicity

**DOI:** 10.1101/620401

**Authors:** Xue Yang, Jonathan K. Williams, Run Yan, M. Maral Mouradian, Jean Baum

**Affiliations:** Department of Chemistry and Chemical Biology, Rutgers University, Piscataway, New Jersey 08854; RWJMS Institute for Neurological Therapeutics, Rutgers Biomedical and Health Sciences, and Department of Neurology, Robert Wood Johnson Medical School, Rutgers University, Piscataway, New Jersey, 08854

## Abstract

Alpha-synuclein (αS) fibrils are toxic to cells and contribute to the pathogenesis and progression of Parkinson’s disease and other synucleinopathies. β-Synuclein (βS), which co-localizes with αS, has been shown to provide a neuroprotective effect, but the molecular mechanism by which this occurs remains elusive. Here we show that αS fibrils formed in the presence of βS are less cytotoxic, exhibit reduced cell seeding capacity and are more resistant to fibril shedding compared to αS fibrils alone. Using solid-state NMR, we found that the overall structure of the core of αS fibrils when co-incubated with βS is minimally perturbed, however, the dynamics of Lys and Thr residues, located primarily in the imperfect KTKEGV repeats of the αS N-terminus, are increased. Our results suggest that amyloid fibril dynamics may play a key role in modulating toxicity and seeding. Thus, enhancing the dynamics of amyloid fibrils may be a strategy for future therapeutic targeting of neurodegenerative diseases.

## 1. Introduction

Parkinson’s disease (PD) is a progressively debilitating neurodegenerative disorder that is estimated to affect 1% of the world’s population over the age of 60^1^. Amyloid fibril deposits of the protein alpha-synuclein (αS) are found in Lewy bodies (LB) and Lewy neurites (LN)^2,3^ in the substantia nigra and other brain regions of PD patients. Myriad evidence shows that αS fibrils are toxic to cells^4–7^, yet the precise role of αS in the pathology of PD and other synucleinopathies is still unclear.

Several hypotheses have been proposed to explain the experimentally observed cellular toxicity of the fibrils. Among these, seeding-propagation is a proposed mechanism to explain the experimentally observed cytotoxicity of the fibrils and the progressive nature of the disease. This process involves the release of mature αS amyloid seeds from the cell^8–10^ that can then be taken up by a neighboring cell^10,11^; these seeds then template the further misfolding and aggregation of endogenous monomeric αS in the recipient cell^7,12^. Fibril polymorphism and protofilament packing have been shown to play an important role in seeding capacity and toxicity^6,13–16^, while the ability of the fibrils to “shed” oligomer and protofibril species may also contribute to cellular toxicity and propagation^17,18^. Previous studies have identified some of the cellular-level details of the internalization, seeding, and propagation of αS fibrils^10,11,19–27^. However, these studies lack the information needed to understand the molecular details of how fibrils can template further aggregation, and critically, the mechanisms by which αS fibril seeding of endogenous αS is affected by inhibitors of αS aggregation.

Beta-synuclein (βS), a homologous protein which is co-localized with αS and is expressed at variable levels relative to αS in different synucleinopathies^28,29^, has been recognized as a natural inhibitor of αS aggregation^30^. A transgenic mouse model that simultaneously expresses both human αS and βS had fewer inclusions and less neurodegeneration compared with only αS-expressing transgenic mice^30^. Interestingly, no detectable amount of βS has been found in LB^31,32^ even though βS can be over-expressed in certain parts of the PD brain^28^, begging the question of how exactly βS interacts with αS to provide neuroprotection and influence αS fibril-induced cellular toxicity.

We previously investigated the sequence and domain level interactions that mediate the influence of βS on the aggregation and fibril formation of αS^33,34^. We have found that head-to-tail transient complexes between βS and αS^35^, mediated by multi-pronged N- and C-terminal interactions^33^, provide enough of a kinetic trap at the earliest stages of αS aggregation to slow down the assembly of αS into fibrils. However, even though βS slows down αS aggregation and reduces the overall αS fibril load in a concentration dependent manner^35,36^, it does not fully abolish αS fibril formation. A detailed understanding of the conformational properties and cytotoxicity of αS fibrils formed in the presence of βS will provide us with a deeper understanding of the mechanisms underlying fibril toxicity.

Here, we show that co-incubation of the monomeric intrinsically disordered αS with the monomeric intrinsically disordered βS results in “co-incubated αS/βS” fibrils that show a significant reduction in cellular toxicity, reduction in seeding capacity and are more resistant to fibril shedding. Solid-state NMR experiments revealed that while the overall structure of the core of αS/βS fibrils is minimally perturbed, the imperfect KTKEGV consensus motif repeats of αS in the N-terminus become dynamic and more water accessible. Our results offer insight into the mechanism of amyloid fibril toxicity and highlight that increased dynamics of co-incubated αS/βS fibrils may interfere with their templating ability, thereby reducing their seeding capacity. Targeting amyloid fibrils by enhancing their dynamics may be a new strategy in designing therapeutics against neurodegenerative diseases.

## 2. Results

### Co-incubation with βS Induces Subtle Differences in αS Fibril Morphology

We studied the differences in the morphology of αS fibrils formed from the incubation of monomeric N-terminally acetylated αS, and αS fibrils formed by co-incubation of monomeric N-terminally acetylated αS with monomeric N-terminally acetylated βS at a 1:3 ratio (called αS/βS co-incubated fibrils). Consistent with our previous work, βS delays αS fibril formation in the Thioflavin T (ThT) aggregation assay (**Fig. S1**), whereby the co-incubation of αS with βS results in a longer lag time and slower growth kinetics compared with αS by itself. Fibrils formed as the end products of these two monomer aggregation assays display slightly different morphologies. Atomic force microscopy (AFM) images show that αS by itself (**Fig. 1a**) forms long straight or twisted fibrils, similar to previous reports^13^, while co-incubated αS/βS forms fibrils that are shorter and straight, with no discernable twisting pattern. On average, the height of αS fibrils (6.0 ± 1.1 nm) tends to be shorter than αS/βS fibrils (7.9 ± 1.7 nm) (**Fig. 1c**), while the length of αS/βS fibrils (0.3 ± 0.2 μm) tend to be shorter than αS fibrils (0.5 ± 0.4 μm) (**Fig. 1d**).

**Figure 1.**
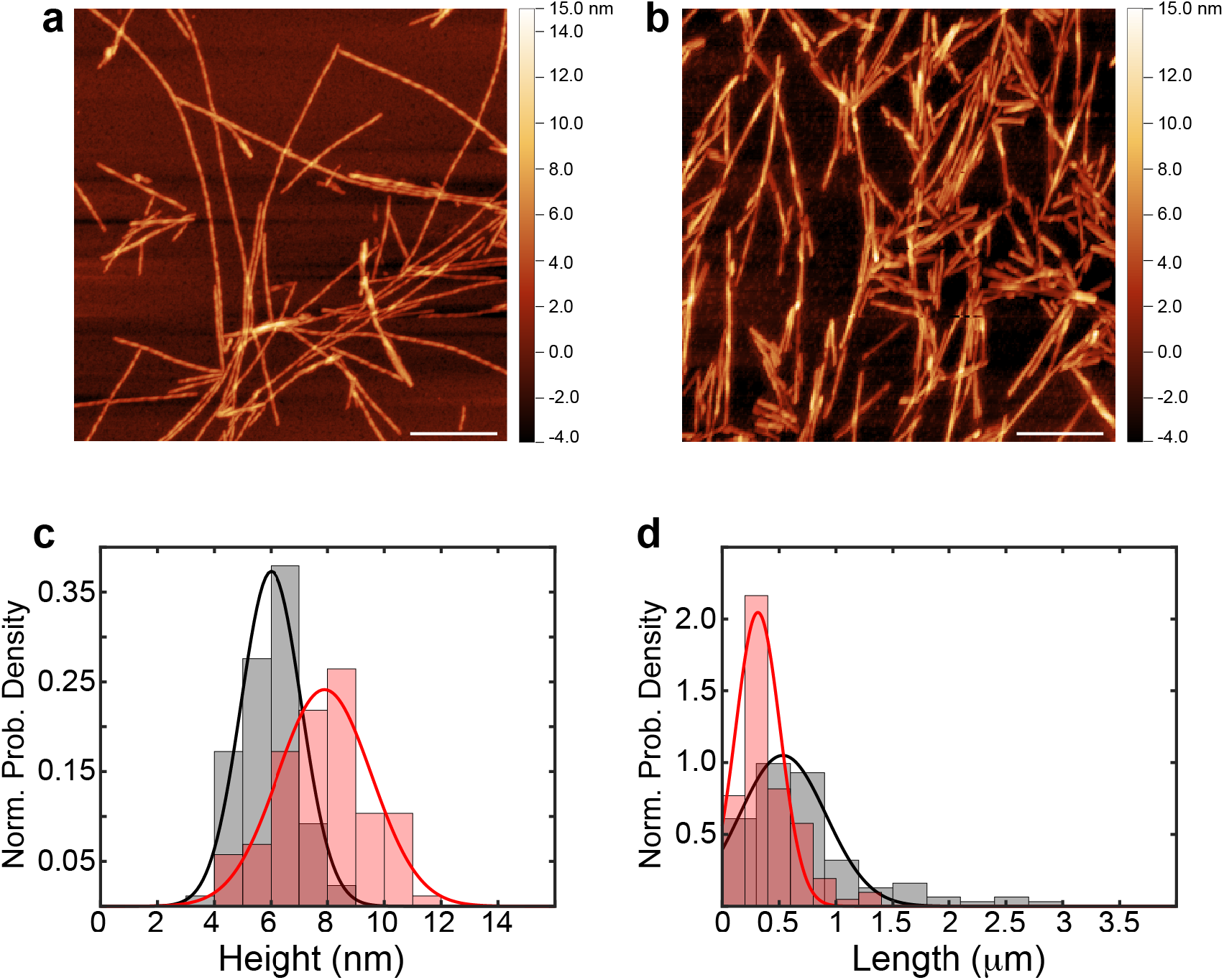
Morphological differences between αS and co-incubated αS/βS fibrils. **(a,b)** AFM images of fibrils formed from 70 μM monomeric αS in **(a)**, and 70 μM monomeric αS plus 210 μM monomeric βS in **(b)**. The length scale bar is 500 nm. Assessment of the height **(c)** and length **(d)** of αS (black) versus co-incubated αS/βS (red) fibrils. Histograms of height and length data are presented as normalized probability densities, and the best-fit probability density function is overlaid.

These AFM images indicate that the primary morphological distinctions between αS and αS/βS fibrils lie in the average height and length, although these parameters are widely distributed (**Fig. 1c,d**). A more subtle distinction lies with the twist of the fibrils: αS fibrils show twisted morphologies (**Fig. 1a**) while αS/βS fibrils do not (**Fig. 1b**). We quantified the monomer composition of the co-incubated αS/βS fibrils to ascertain whether these morphological changes were induced by incorporation of βS into the fibril body. Mature fibrils were solubilized in 4M guanidine hydrochloride and analyzed by ESI-MS. Surprisingly, the co-incubated αS/βS fibrils are composed of less than 6% βS (**Fig. S1**). These observations suggest that βS is more likely surface associated with αS fibrils, rather than interleaved within the body of the fibril, supported by the larger height profiles, shorter length profiles, and the fact that very little βS is retained in mature αS/βS fibrils.

### αS/βS co-Incubated Fibril Core Structure is Minimally Perturbed while the N-Terminal Dynamics are Increased

We investigated the conformational and dynamics properties of the αS fibril when coincubated with βS utilizing solid-state NMR (ssNMR) spectroscopy. **Figure 2** shows the water-edited 2D ^13^C-^13^C correlation spectra^37,38^ of ^13^C,^15^N-labeled αS fibrils (**Fig. 2a**) and co-incubated αS/βS fibrils where only αS is uniformly labeled with ^13^C and ^15^N (**Fig. 2b**). This experiment utilizes a cross-polarization period to transfer ^1^H to ^13^C magnetization, which preferentially detects the rigid residues that make up the core of the fibril and does not detect the dynamic or disordered residues that make up the bulk of the N- and C-terminal regions. The long water ^1^H spin-diffusion (SD) mixing time (100 ms, black) spectra represent a state where the water ^1^H magnetization has fully equilibrated across the fibril, while the short water ^1^H SD mixing time (3 ms, red) spectra illustrate the fibril residues that are in closest proximity to water. The relative proximity or accessibility of a residue to water is then most easily compared by taking the ratio between these two intensities (Int_3ms_/Int_100ms_).

**Figure 2.**
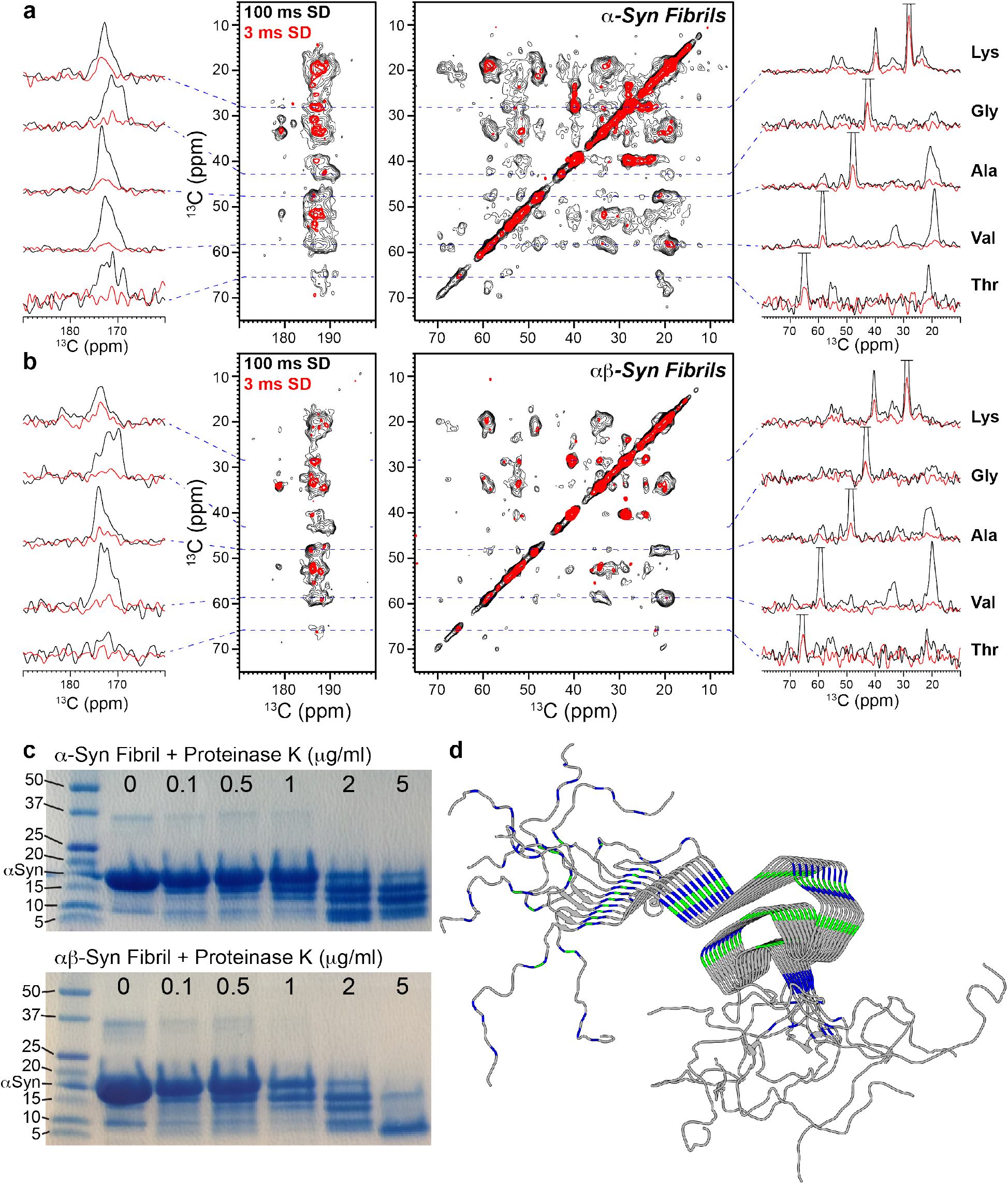
Changes in fibril water accessibility and fibril degradation. Water-edited solid-state NMR ^13^C-^13^C correlation spectra of **(a)** αS and **(b)** co-incubated αS/βS fibrils. On either side of the 2D spectra are 1D slices of the carbonyl region (left side) and aliphatic region (right side), showing the intensities of cross-peaks to several amino acid types. Magnetization has equilibrated at long water spin-diffusion times (100 ms, black) compared with the initial water-protein magnetization transfer at short spin-diffusion times (3 ms, red). The ratio of these intensities (Int_3ms_/Int_100ms_) indicates the relative proximity of water to the various residues in the protein. **(c)** Digestion of αS and αS/βS fibrils at various concentrations of proteinase K. Full-length gels are presented in Supplementary Figure 6. **(d)** Map (PDB ID: 2N0A) of the residues that show the largest degree of change in water accessibility between αS and αS/βS fibrils, lysine (blue) and threonine (green).

The ^13^C chemical shifts of the long SD mixing time spectra do not show marked differences between αS and αS/βS fibrils, indicating that the core structure of the fibril does not change even when formed in the presence of a stoichiometric excess of βS (**Fig. 2a,b**). However, there are some subtle differences in peak intensities and water-fibril SD buildup between αS and αS/βS fibrils. The side chain Cβ-Cα and Cβ-Cγ cross-peaks of the threonine residues are well resolved from chemical shift overlap of any other residue in the region from ~65-70 ppm (**Fig. 2**). Slices from the water edited 2D ^13^C-^13^C spectra show decreases in the relative intensities of several of the threonine cross-peaks of the αS/βS fibrils compared to αS fibrils, while those threonine cross-peaks that remain in the αS/βS fibril spectra have increased water spin-diffusion (i.e. larger Int_3ms_/Int_100ms_ ratios) relative to αS fibrils (**Fig. 2**). The loss of intensity of αS/βS fibril threonine peaks could be caused by increased dynamics of these residues, while the increase in water spin-diffusion of the αS/βS fibril threonine peaks indicates that these residues are more water accessible. The side chain Cδ cross-peaks of the lysine residues resonate around ~29 ppm, and also show increases in water spin-diffusion ratios of αS/βS fibrils relative to αS fibrils (**Fig. 2**). Conversely, the water spin diffusion ratios of the hydrophobic alanine and valine residues do not change between αS and co-incubated αS/βS fibrils (**Fig. 2a,b**), indicating that the hydration environment of these residues does not significantly change between the two fibrils.

αS has a 140 amino acid primary sequence generally described by 3 domains: a 60 residue polyampholyte N-terminal domain, a 35 residue hydrophobic NAC domain, and a 45 residue highly negatively-charged polyelectrolyte C-terminal domain. Lysine and threonine (**Fig. 2d**) are almost exclusively located in the N-terminal and NAC regions of the αS sequence, where they are clustered in the imperfect KTKEGV repeats. These two regions form the “Greek-key” motif of the αS fibril core structure, suggesting that βS influences the αS fibril by increasing the dynamics of these regions.

### Co-Incubated Fibrils are More Sensitive to Proteinase K Digestion

Proteasomal impairment has been implicated in several neurodegenerative diseases^39^, including PD^40^, and as proteasome activity decreases with age cells become more vulnerable to deleterious protein aggregation^41^. Therefore, an understanding of how synuclein fibrils and aggregates undergo protease degradation and clearance may shed critical light on PD progression. In order to understand the differences in protease degradation and fibril stability between αS fibrils and co-incubated αS/βS fibrils, we carried out a series of digestion assays with increasing concentrations of proteinase K (**Fig. 2c**). We observed that co-incubated αS/βS fibrils are more sensitive to proteinase K digestion, displaying an increased fibril digestion profile at lower concentrations of proteinase K (1 μg/ml) relative to αS fibrils (2 μg/ml). In addition, αS/βS fibril degradation products are largely small molecular species (< ~5 kDa), whereas αS fibrils show a range of degradation products (~5-15 kDa) at 5 μg/ml proteinase K concentration.

These digestion profiles indicate that αS/βS fibrils are more easily accessible to cleavage by proteinase K, suggesting that the co-incubated αS/βS fibrils might be more susceptible to degradation in vivo (**Fig. 2c**). These profiles resemble those from previous work by Miake and coworkers, who established that proteinase K digestion of αS preferentially cleaves the N- and C-terminal portions of αS and leaves the fibril core from residues 31-109 intact^42^. Taken together with the ESI-MS and ssNMR data, these results again suggest that βS is surface associated along αS fibrils and induces more dynamic flexibility in portions of the αS fibril (i.e. the N-terminal and C-terminal domains).

### Co-Incubated αS/βS Fibrils are Less Toxic and Exhibit Reduced Seeding and Proliferation Capacity Compared to αS Fibrils in Neuroblastoma Cells

We investigated how co-incubating βS with αS affects the cytotoxicity of the fibrils. Fibrils of αS or αS/βS were added to cultures of human SH-SY5Y neuroblastoma cells and incubated for 24 hours, at which time cellular viability was assessed by the ability of the cells to reduce 3-(4, 5-dimethylthiazol-2-yl)-5-(3-carboxymethoxyphenol)-2-(4-sulfophenyl)-2H-tetrazolium (MTS). Compared with untreated cells, or cells treated with monomeric αS, fibrils of αS induced a 30% reduction in cell viability (***p < 0.001), whereas αS/βS fibrils had no impact on cell viability (**Fig. 3a**).

**Figure 3.**
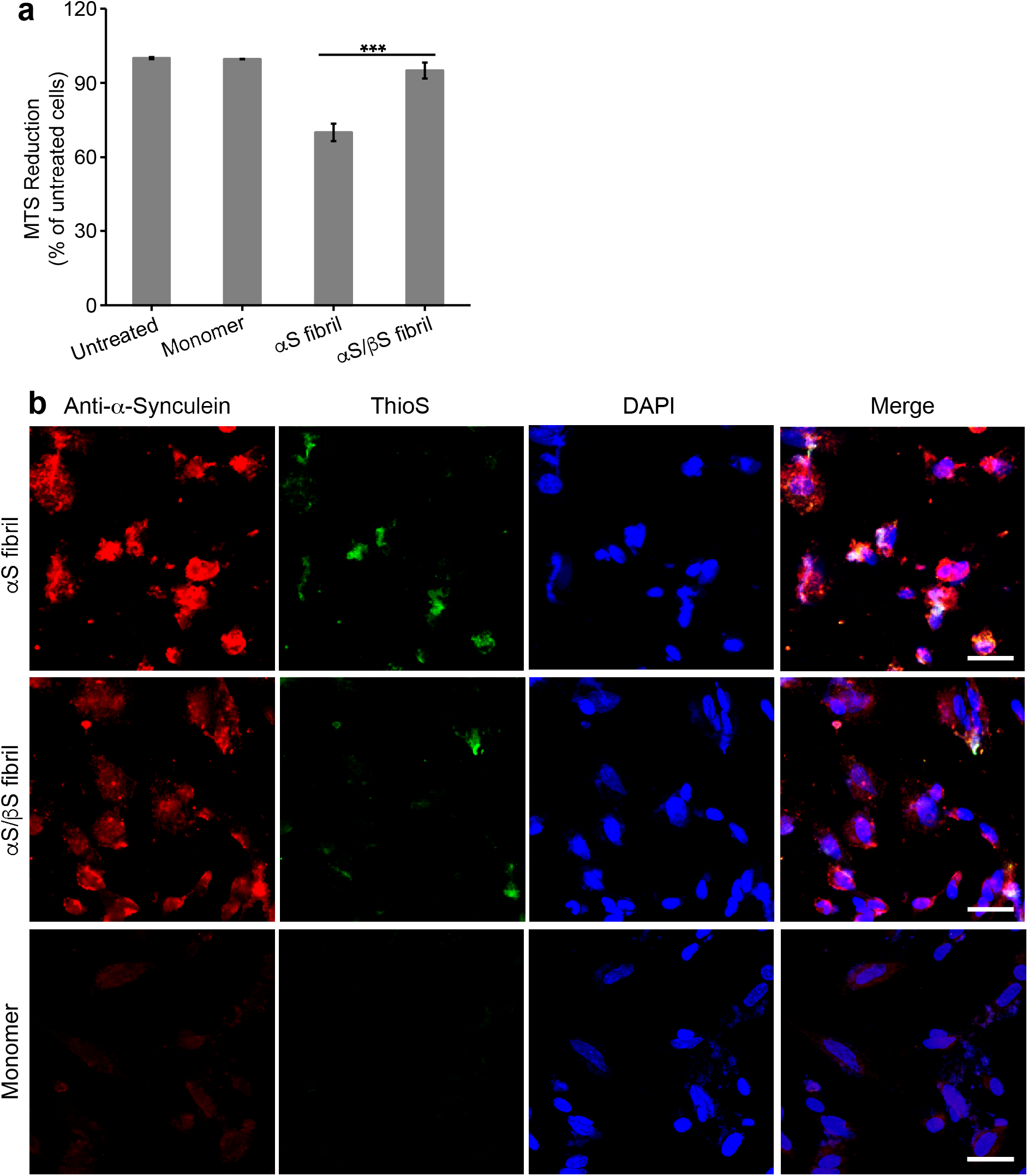
Cellular toxicity of αS and co-incubated αS/βS fibrils and their seeding potential. **(a)** Viability of SH-SY5Y cells assessed by MTS assay after treatment with αS or αS/βS fibrils (1.3 μM monomer equivalents), or monomeric αS (1.3 μM) as control, for 24 hours. Data shown are means and standard errors of the mean (SEM) of 3 independent experiments run in triplicates. ***ANOVA p < 0.001. **(b)** Confocal microscopy images of SH-SY5Y cells treated with 1.3 μM αS fibril (top), αS/βS fibril (middle), or αS monomer (bottom) for 24 hours before fixing and staining. Separate channels are presented showing the presence of all αS species (anti-αS antibody, red), all amyloid species (ThioS, green), and cell nuclei (DAPI, blue), along with the three channels overlaid (merge). The scale bar is 40 μm.

We also investigated the ability of αS and αS/βS fibrils to seed further aggregation of endogenous αS in SH-SY5Y cells by analyzing the fluorescence intensities of dyes that specifically bind to αS and amyloid structures (**Fig. 3b**). Cells were treated with monomeric αS, αS fibrils or αS/βS fibrils for 24 hours before being fixed and stained with purified mouse anti-αS (anti-α-synuclein) antibody, thioflavin S (ThioS), and 4’,6-diamidino-2-phenylindole (DAPI). Cells were then imaged by confocal fluorescence microscopy, where the anti-αS antibody fluoresces red and indicates the presence of any synuclein species present, ThioS fluoresces green and indicates the formation of amyloid species, and DAPI stains the cell nucleus blue (**Fig. 3b**). Compared with cells treated with monomeric αS (**Fig. 3b**, bottom row), cells treated with αS fibrils showed an increase in anti-αS antibody fluorescence of 7.3x (**Fig. 3b**, top row), while cells treated with αS/βS fibrils showed a smaller increase of 4.4x (**Fig. 3b**, middle row). ThioS staining indicating amyloid formation showed a similar trend with a 4.8x increase with αS fibrils vs a 3.4x increase with αS/βS fibrils.

These results demonstrate that while αS fibrils are indeed toxic to neuroblastoma cells (**Fig. 3a**), co-incubated αS/βS fibrils are not (**Fig. 3a**). In addition, co-incubated αS/βS fibrils have a lesser tendency to cause the formation of synuclein aggregates and amyloid species compared with αS fibrils (**Fig. 3b**). Taken together, co-incubation of βS with αS results in fibrils that are not toxic to cells and have reduced ability to seed further aggregation in a cellular environment. The reduced seeding ability also suggests that βS interferes with the ability of αS fibrils to catalyze secondary nucleation processes on the fibril surface, indicating that βS is likely surface associated along the αS fibril.

### Oligomers Shed from αS or αS/βS Fibrils Have Different Morphologies, Toxicities and Seeding Capacities

It has been hypothesized that as the endpoint of misfolding and aggregation of several neurodegenerative disease associated proteins, amyloid fibrils might act as a “sink” to sequester misfolded toxic species^43^. However, amyloid fibrils do not represent a completely stable species in solution, rather they exist in a dynamic equilibrium between fibril and oligomer forms. Indeed, toxic oligomers have even been observed to shed from mature αS fibrils over time^18^. To understand the effect of βS on the stability and equilibrium of αS fibrils, we sought to determine the morphology, toxicity and cell seeding capacities of the oligomers that are shed from αS fibrils and αS/βS fibrils.

We first measured the thermostability of the two fibrils using far-UV circular dichroism (CD) spectroscopy. The CD spectra show that both αS and αS/βS fibrils have the characteristic spectral minimum at 218 nm, indicating the presence of β-sheet structure (**Fig. S2**). We monitored the change in ellipticity of the 218 nm signal as a function of temperature, and found that change in ellipticity of co-incubated αS/βS fibrils is less than that of αS fibrils as temperature increased, indicating that αS/βS fibrils are more thermostable than αS fibrils (**Fig. S2**). AFM images show that the oligomers that are shed from αS fibrils (**Fig. 4a**) primarily adopt small globular morphologies, while oligomers shed from αS/βS fibrils tend to adopt short protofibril morphologies with some larger globular species also present (**Fig. 4b**). We next measured the toxicity of the shed oligomers in SH-SY5Y cells. After a 48 hour period of incubation with shed oligomers from either αS or αS/βS fibrils, we found that oligomers shed from αS reduced cell viability by 17% compared to the untreated cells and cells treated with monomeric αS, whereas oligomers shed from αS/βS did not (**Fig. 4c**). We also assessed the ability of shed oligomers to seed further aggregation in cells, using confocal fluorescence microscopy. Compared with cells treated with monomeric αS (**Fig. 4d**, bottom row), cells treated with oligomers shed from αS fibrils showed an increase in anti-synuclein antibody fluorescence of 1.6x (**Fig. 4d**, top row), while cells treated with oligomers shed from αS/βS fibrils showed an increase of 1.3x (**Fig. 4d**, middle row). ThioS staining indicates that amyloid formation increased by 1.6x in cells treated with oligomers shed from αS fibrils and by 1.3x in cells treated with oligomers shed from αS/βS fibrils.

**Figure 4.**
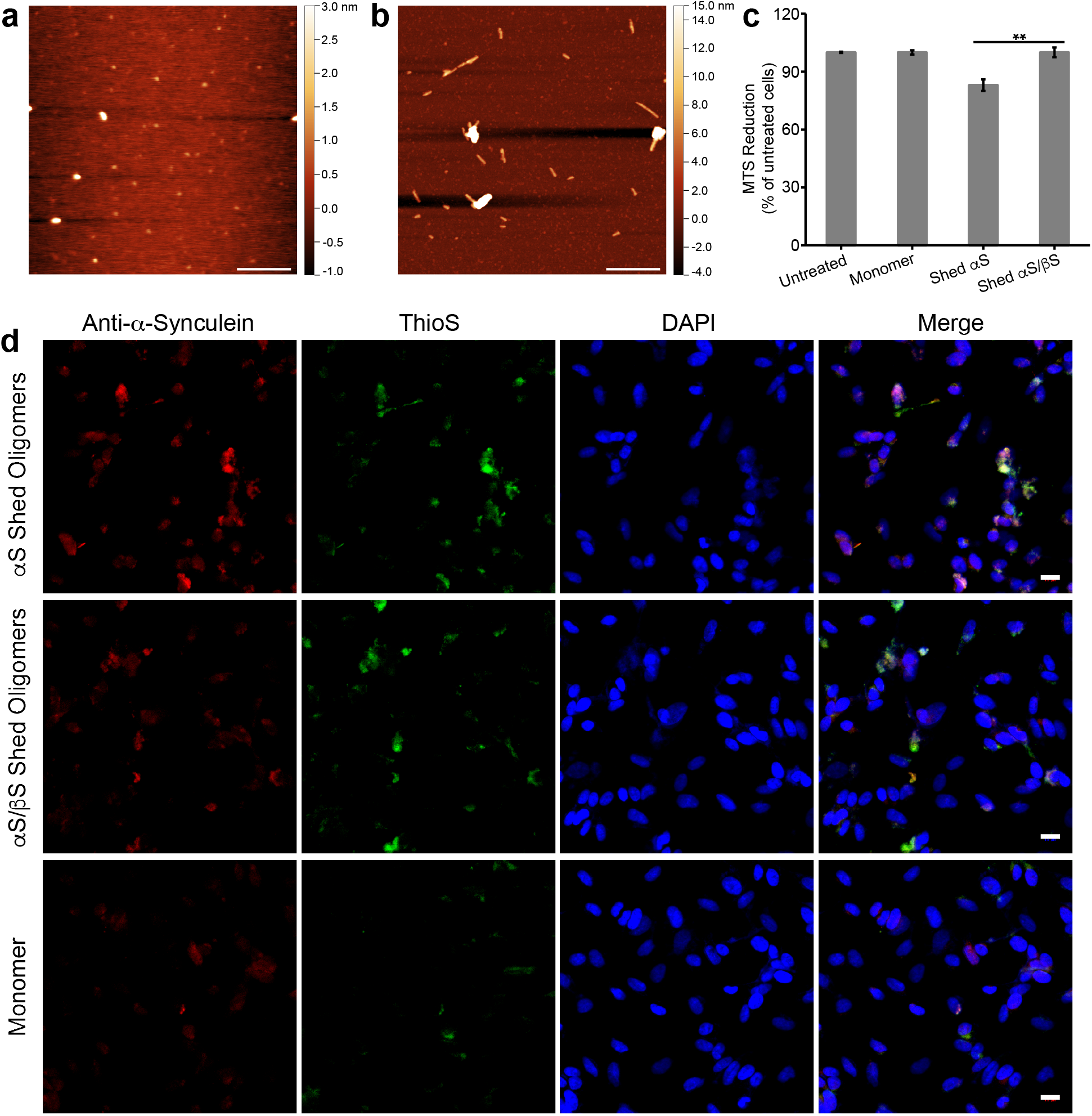
Morphology and toxicity of oligomeric species that are shed from mature fibrils. AFM images of the oligomeric species that are shed from mature αS fibrils **(a)** and mature αS/βS fibrils **(b)**. The length scale bar is 500 nm. **(c)** Viability of SH-SY5Y cells assessed by MTS assay after treatment for 48 hours with the shed oligomers from αS fibrils or shed oligomers from αS/βS fibrils (0.7 μM monomer equivalents), or with monomeric αSyn (0.7 μM) as a control. Data shown are means ± SEM of 3 independent experiments run in triplicates. ** ANOVA p < 0.01. **(d)** Confocal microscopy images of SH-SY5Y cells treated with oligomers shed from αS fibrils (top), oligomers shed from αS/βS fibrils (middle), or with αS monomer (bottom) for 48 hours before fixing and staining. Separate channels are presented showing the presence of all αS species (anti-αS antibody, red), all amyloid species (ThioS, green), and cell nuclei (DAPI, blue), along with the three channels overlaid (merge). The scale bar is 16 μm.

**Figure 5.**
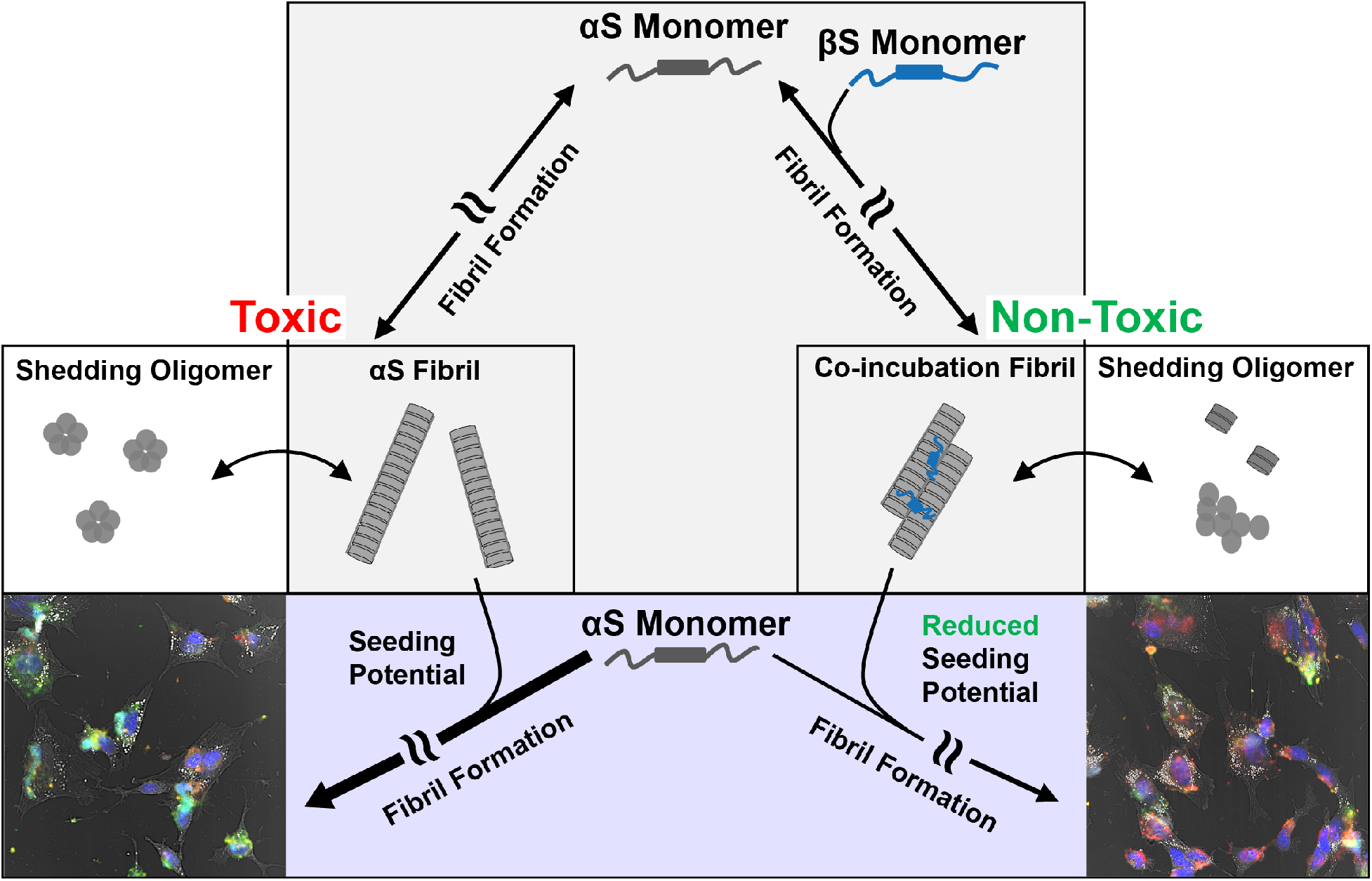
αS and αS/βS fibril toxicities and seeding potentials. αS aggregates and misfolds along a nucleation-dependent fibril formation pathway, generating various oligomeric species before finally adopting a characteristic repeating cross-beta amyloid fibril structure. When αS aggregates on its own, the resulting fibrils are toxic to cultured human neuroblastoma cells (*left pathway*), and the oligomeric species that shed from these fibrils are also toxic to cells. However, if αS is co-incubated with βS and allowed to aggregate, then the resulting fibrils are no longer toxic to cells (*right pathway*), and oligomer species that shed from these fibrils are also nontoxic. These different fibrils also display differential seeding capacities (*confocal images, bottom*). αS fibrils are able to efficiently seed amyloid formation, while co-incubated αS/βS fibrils have reduced propensity to seed further aggregation, as evidenced by the difference in green intensity in confocal images.

Our observations show that αS fibrils are less thermostable (**Fig. S2**) and shed primarily small globular and amorphous oligomers (**Fig. 4a**), while co-incubated αS/βS fibrils are more thermostable (**Fig. S2**) and shed primarily short proto-fibril aggregates (**Fig. 4b**). The proto-fibril species shed from αS/βS fibrils also show reduced seeding propensity compared to the small globular and amorphous αS oligomer species (**Fig. 4d**). This finding suggests that the dynamic equilibrium is shifted away from the formation of small toxic oligomers towards less toxic proto-fibrils in the presence of βS (**Fig. 4c**). Taken together in the context of our previous observations, βS appears to interact with the unstructured regions of αS fibrils to stabilize the core structure of co-incubated αS/βS fibrils, while increasing the local dynamics of the N-terminal domain. This results in a reduced capacity for seeding and shedding of toxic oligomeric species.

## 3. Discussion

Amyloid fibrils of αS are key pathologic features of PD and have been recognized as contributing to the progression of the disease. These fibrils are thought to contribute to cellular toxicity through their ability to seed further aggregation of endogenous αS, and the ability of the fibrils to “shed” oligomer and protofibril species that may be toxic.The αS fibrils studied in this work should be distinguished from fibrils that are contained within aggresome-like LBs. Fibrils that are formed as the end product of the aggregation pathway of αS (either *in vivo* or *in vitro*), and are not yet collected into LBs, exist in a dynamic equilibrium with oligomers, as evidenced by the ability of fibrils to “shed” smaller molecular species^18,44^. Here we have demonstrated that αS fibrils formed in the presence of the natural inhibitor βS, while maintaining similar core structures as αS fibrils alone, exhibit reduced toxicity to neuroblastoma cells, reduced seeding properties, and are in dynamic equilibrium with oligomers that also share reduced toxicity and seeding.

We have utilized ssNMR and the changes in ^13^C chemical shifts to probe how the core residues and dynamics of αS fibrils and co-incubated αS/βS fibrils differ from one another. While we have not yet completed a full assignment and structure determination of our fibrils, ^13^C chemical shifts are very sensitive reporters of amino acid type and secondary structure^47^. The ^13^C resonances observed in our spectra show characteristic β-sheet chemical shifts. Comparison of our 1D ^13^C, 2D ^13^C-^13^C and 2D ^15^N-^13^C (**Fig. S3-5**) spectra with the chemical shift lists (BMRB Entry 25518) and spectra reported by Rienstra and coworkers^48^, who have previously determined the full-length αS fibril structure by ssNMR, show relatively good agreement. Since we have prepared our fibrils in a similar manner to those used for the published αS fibril structure^48^, we can reasonably assume that our fibrils adopt a similar structure. Previously, residues 55-62 were identified as being disordered in the αS fibril^48^. Based on the assignments of the threonine and lysine cross-peaks in our 2D ^13^C-^13^C spectra of the co-incubated αS/βS fibril, we observe that water accessibility increases, and local residue dynamics arise, in the imperfect repeat KTKEGV consensus motifs that are found in the N-terminal and NAC domains of co-incubated αS/βS fibrils relative to αS fibrils alone.

βS has previously been identified in studies of transgenic mice as a natural anti-Parkinsonian factor which has the ability to reduce αS inclusion formation^30^. Yet, even though it reduces αS positive inclusions, it does not completely abolish the formation of αS fibrils. We propose that the role of βS as an inhibitor is multifaceted, influencing αS aggregation at multiple points along its fibril-formation pathway. In the earliest stages of αS aggregation, βS can stabilize αS in αS-βS heterodimers^35^, which help to slow down the conversion of αS into higher order aggregates. As αS continues to aggregate, βS has been found to stabilize and eliminate the formation of toxic oligomers^45,46^. In this work, we have now shown that in the last stage of αS aggregation, co-incubation with βS minimizes the toxicity and seeding ability of αS fibrils, and furthermore alters the fibril-oligomer equilibrium. Our findings demonstrate that βS can reduce the effects of toxic αS fibrils in cells without changing the core structure of αS fibrils, and provide insight into how the dynamics and the surface of these fibrils may directly contribute to their toxicity and seeding ability. The multi-pronged targeting of αS by βS highlights the potential of βS as a lead for the future design of inhibitors that provide therapeutic intervention in synucleinopathies at multiple stages of αS aggregation.

The misfolding and aggregation of endogenous αS monomers due to seeding by fibrils is believed to be critical to the progression of synucleinopathies. The mechanism by which mature αS fibrils seed further aggregation is believed to proceed by surface-mediated secondary nucleation^49–51^, where the surface properties of αS fibrils govern their interaction with endogenous αS monomers and template further aggregation. The exact details of how additional αS monomers undergo templated conversion are not yet known, but the present work provides some clues. The recent cryo-EM structures of αS fibrils show that a steric-zipper motif in the N-terminal domain mediates the interface between two protofilaments and stabilizes the mature fibril morphology^13,52^. Our results show that co-incubated αS/βS fibrils have increased dynamics and water accessibility of residues in the N-terminal domain and allow for enhanced protease degradation of the fibril, suggesting that the protofilament interface may be altered and more dynamic. These observations also highlight the importance of dynamics in mediating the seeding ability of αS fibrils: increased dynamics of the N-terminal domain may lead to reduced seeding, as secondary nucleation may necessitate a rigid N-terminal domain for proper templating of αS aggregation. Our results suggest that enhancing amyloid fibril dynamics at templating domains may be an approach for future therapeutic intervention for neurodegenerative diseases.

## 4. Materials and Methods

### Protein Expression and Purification

Expression of N-terminally acetylated human αS and βS proteins was performed via co-expression with pNatB plasmid (Addgene #53613) in *E. coli* BL21(DE3) cells, and protein purification was performed as described previously^53^. Uniformly ^13^C, ^15^N isotopically labeled αS for ssNMR experiments was expressed in M9 minimal media supplemented with ^13^C-glucose and ^15^N-ammonium chloride as the sole carbon and nitrogen sources, respectively. Protein molecular weight and purity were assessed by ESI-MS, and stored at −20 °C as a lyophilized powder until use.

### Fibril Sample Preparation

Lyophilized acetylated αS or βS was dissolved in 10 mM PBS (pH 7.4), and large aggregates were removed by centrifuge filtration (50 kDa MWCO, Millipore Sigma, St. Louis, MO). The dissolved protein was concentrated in 3 kDa centrifuge units (Millipore Sigma, St. Louis, MO) to 1 mg/mL (αS) or 3 mg/mL (βS). To create fibrils, 100 uL of each sample mixture was loaded into 96-well clear bottom plates (Corning, Corning, NY) with a single Teflon bead (3 mm, Saint-Gobain N.A., Malvern PA). The plates were sealed with Axygen sealing tape (Corning, Corning, NY) and shaken at 600 rpm and 37 °C in a POLARstar Omega fluorimeter (BMG Labtech, Cary, NC). Fibrils were allowed to form for at least 72 hours. Samples used for AFM, ESI-MS, ssNMR, and cell toxicity and shedding experiments were collected by centrifugation at 14k rpm for 2 hours, and washed through multiple rounds of re-suspension in 10 mM PBS (pH 7.4) and centrifugation at 14k rpm for 2 hours in order to remove residual soluble and non-fibrillar components.

### Preparation of Oligomer Species Shed from Fibrils

Fibril samples were re-suspended in 1 mL of 10 mM PBS (pH 7.4) and incubated at 37 °C for 72 h, followed by removal of mature fibrils by using 0.22 μm filter (Millipore Sigma, St. Louis, MO). Samples were concentrated with 3 kDa centrifuge units (Millipore Sigma, St. Louis, MO), and protein concentration was measured using a bicinchoninic acid (BCA) assay (Thermo Scientific, Waltham, MA).

### Proteinase K Digestion

Fibrils at a concentration of 1 mg/mL were incubated with various concentrations (0.1, 0.5, 1.0, 2.0, 5.0 μg/mL) of proteinase K (Sigma Aldrich, St. Louis, MO) in 10 mM PBS (pH 7.4) at 37 °C for 1 h. The digestion reaction was quenched by the addition of a 1200:1 molar excess of phenylmethane sulfonyl fluoride (Sigma Aldrich, St. Louis, MO) followed by the addition of 2 M guanidine thiocyanate (Sigma Aldrich, St. Louis, MO) and incubation at room temperature for 4 h. The results of the degradation reaction were mixed with 4x SDS-PAGE loading buffer (Invitrogen, Carlsbad, CA), loaded onto precast gels (Bio-Rad, Hercules, CA), and run at 120 V for 50 min.

### Atomic Force Microscopy (AFM)

Samples (20 μL) were placed onto freshly cleaved mica (Ted Pella Inc., Redding, CA) and incubated for 15 min at room temperature, followed by 3 washes of 200 μL each deionized water as described previously^34^. All images were collected on a NX-10 instrument (Park Systems, Suwon, South Korea) using non-contact mode tips (PPP-NHCR, 42 N/m, 330 kHz; Nanosensors, Neuchatel, Switzerland). Image processing and analysis were carried out in the Gwyddion software package^54^.

### Solid-State Nuclear Magnetic Resonance Experiments

All MAS ssNMR experiments were carried out on an Avance III HD 600 MHz (14 T) spectrometer (Bruker BioSpin, Billerica, MA) using a 1.6 mm triple resonance MAS probe (Phoenix NMR, Loveland, CO) tuned to ^1^H/^13^C/^15^N frequencies. Typical radiofrequency (rf) field strengths were 118 kHz for ^13^C, 74 kHz for ^15^N, and 100-145 kHz for ^1^H. ^13^C chemical shifts were referenced to the ^13^CH_2_ signal of adamantane at 38.48 ppm on the tetramethylsilane scale, and ^15^N chemical shifts were referenced to the ^15^N signal of N-acetylvaline at 122.0 ppm on the liquid ammonia scale. All experiments utilized a MAS rate of 13.333 kHz, and sample temperature was controlled to 25 °C, unless otherwise noted. One-dimensional (1D) ^13^C MAS spectra were recorded using a conventional cross-polarization (CP) sequence. Two-dimensional (2D) ^13^C-^13^C dipolar-assisted rotational-resonance (DARR) experiments^55^ utilized a mixing period of 100 ms. A 2D water-edited DARR^37^ experiment, with a DARR mixing period of 100 ms, a T_2_-filter of 6 ms, and a ^1^H spin-diffusion period of either 3 ms or 100 ms, was used to measure the water-protein ^1^H spin diffusion differences between the two fibril samples. 2D ^15^N- ^13^C correlation spectra were measured using a REDOR-based pulse sequence^56^, utilizing a REDOR period of 1.35 ms to observe long range correlations.

### Analysis of Fibril Composition by ESI-MS

Mature fibril samples were dissolved in 4 M guanidine hydrochloride overnight, then buffer exchanged with 50 mM ammonium acetate with 0.1% formic acid. Samples were concentrated to 10 μM for ESI-MS analysis.

### Neuroblastoma Cell Culture

Human SH-SY5Y neuroblastoma cells (ATCC, Manassas, VA) were cultured in DMEM/F12 (GE Healthcare, Boston, MA) with 10% fetal bovine serum (Gibco Co., Dublin, Ireland) and kept in a 37 °C, 5% CO_2_ humidified atmosphere. Before cell viability assays or immunocytochemistry, cells were plated into 96-well (Corning, Corning, NY) or 12-well plates (Cellvis, Mountain View, CA), and allowed to grow for 24 h.

### Cell Viability MTS Reduction Assay

SH-SY5Y cells were treated with 1.3 μM fibril (24 h), 0.7 μM shed species (48 h), or an equivalent concentration of monomer as a control (24 h or 48 h). Cell viability was assessed by adding 20 μL MTS per 100 uL cell culture (Promega, USA) and incubating for 2.5 h at 37 °C, before measuring absorbance at 490 nm.

### Immunocytochemistry

SH-SY5Y cells were treated with 1.3 μM fibril (24 h), 0.7 μM shed species (48 h), or an equivalent concentration of monomer as a control (24 h or 48 h). Cells were fixed with 10% formalin (Sigma Aldrich, St. Louis, MO) and permeabilized with 0.5% Triton in PBS (Sigma Aldrich, St. Louis, MO). Cells were then blocked by incubation with 5% Donkey Serum solution (Sigma Aldrich, St. Louis, MO) for 30 min at 37 °C. Cells were incubated with 0.01% thioflavin S (Acros Organics, Waltham, MA) for 10 min at 37 °C and then washed with PBS, followed by incubation with purified mouse anti-α-synuclein primary antibody (BD Biosciences, Franklin Lakes, NJ; Cat.# 610786, RRID: AB_398107) at 4 °C overnight in the dark. Cells were washed with PBS 3 times and incubated with fluorophore-conjugated secondary antibody TRITC (Sigma Aldrich, St. Louis, MO; Cat.# T5393, RRID: AB_261699) for 1 h, then washed again 3 times with PBS. Cells were incubated with DAPI for 1.5 min at room temperature and then washed with PBS, to visual cell nuclei. All samples were imaged using a Zeiss LSM 780 confocal laser scanning microscope with 20x objective (Zeiss, Oberkochen, Germany), and images were processed using the Fiji distribution of ImageJ^57^.

### Statistical Analysis

All cell viability experiments were performed in triplicate, and each assay was repeated at least three times. Statistical significance was determined by one-way ANOVA with post-hoc Bonferonni analysis (GraphPad Prism).

## Acknowledgements

This work was supported by National Institutes of Health (NIH) Grant GM110577 (J.B.). M.M.M. is the William Dow Lovett Professor of Neurology and is funded by NIH grants NS096032, AT006868, NS073994, and NS101134, the Michael J. Fox Foundation for Parkinson’s Research, and the American Parkinson Disease Association.

## Author Contributions

J.B. and M.M.M. conceived the study. X.Y. expressed and purified protein, and prepared fibril and oligomer samples for all experiments. X.Y. performed the proteinase K assay, AFM, and ESI-MS experiments. X.Y. and R.Y. designed and carried out the SH-SY5Y cell culture, MTS assay, and immunocytochemistry experiments. J.K.W. designed and performed the solid-state NMR experiments. X.Y., J.K.W., M.M.M. and J.B. analyzed the data and wrote the manuscript. All authors reviewed, edited and approved the manuscript.

