## Supplemental Information for "Increased Dynamics of α-Synuclein Fibrils by β-Synuclein Leads to Reduced Seeding and Cytotoxicity"

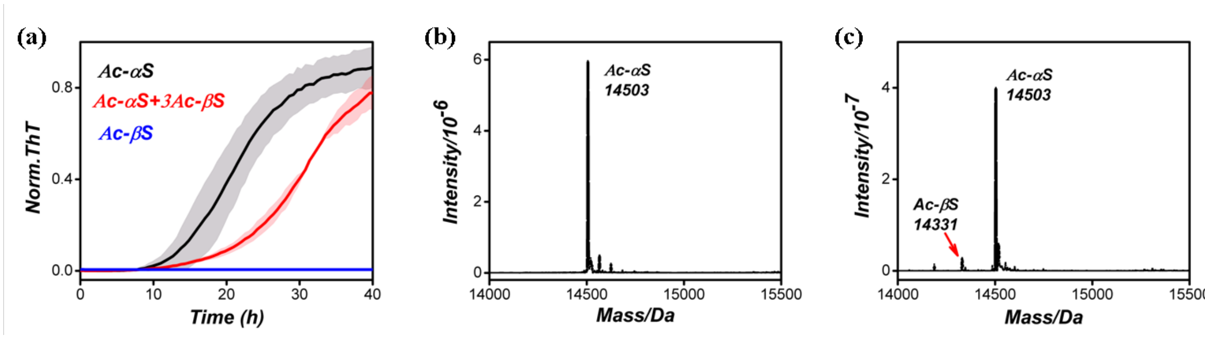


**Figure S1:** (a) Normalized change in ThT fluorescence signal of 70 μM Ac-αS (black), 210 μM Ac-βS (blue), or a mixture of 70 μM Ac-αS + 210 μM Ac-βS (1:3) (red) incubated at 37°C in 10 mM PBS with Teflon beads and shaking. (b,c) ESI-MS data showing the protein composition of αS fibrils and co-incubated αS/βS fibrils.

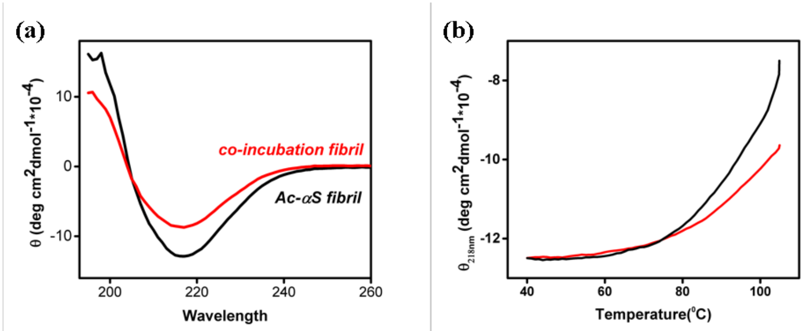


**Figure S2:** (a) Far-UV CD wavelength scan spectrum of αS fibril (black) and co-incubated αS/βS fibril (red) showing similar secondary structure. (b) Thermal stability curve of αS fibril (black) and co-incubation αS/βS fibril (red), monitored by CD signal change at 218nm.

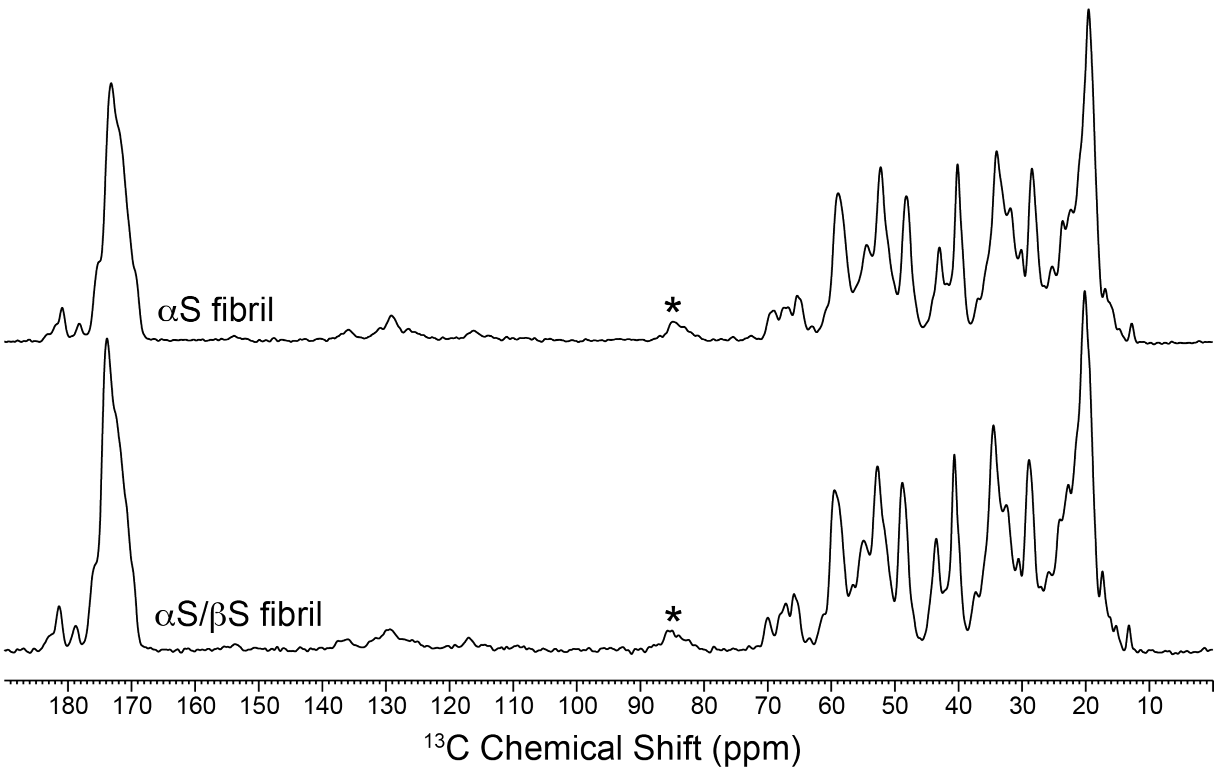


**Figure S3:** One-dimensional (1D) $^{13}C$ cross-polarization (CP) spectra of αS fibrils (top) and co-incubated αS/βS fibrils (bottom). Spectra were recorded at a MAS rate of 13.333 kHz, and temperature was controlled at 25 °C.

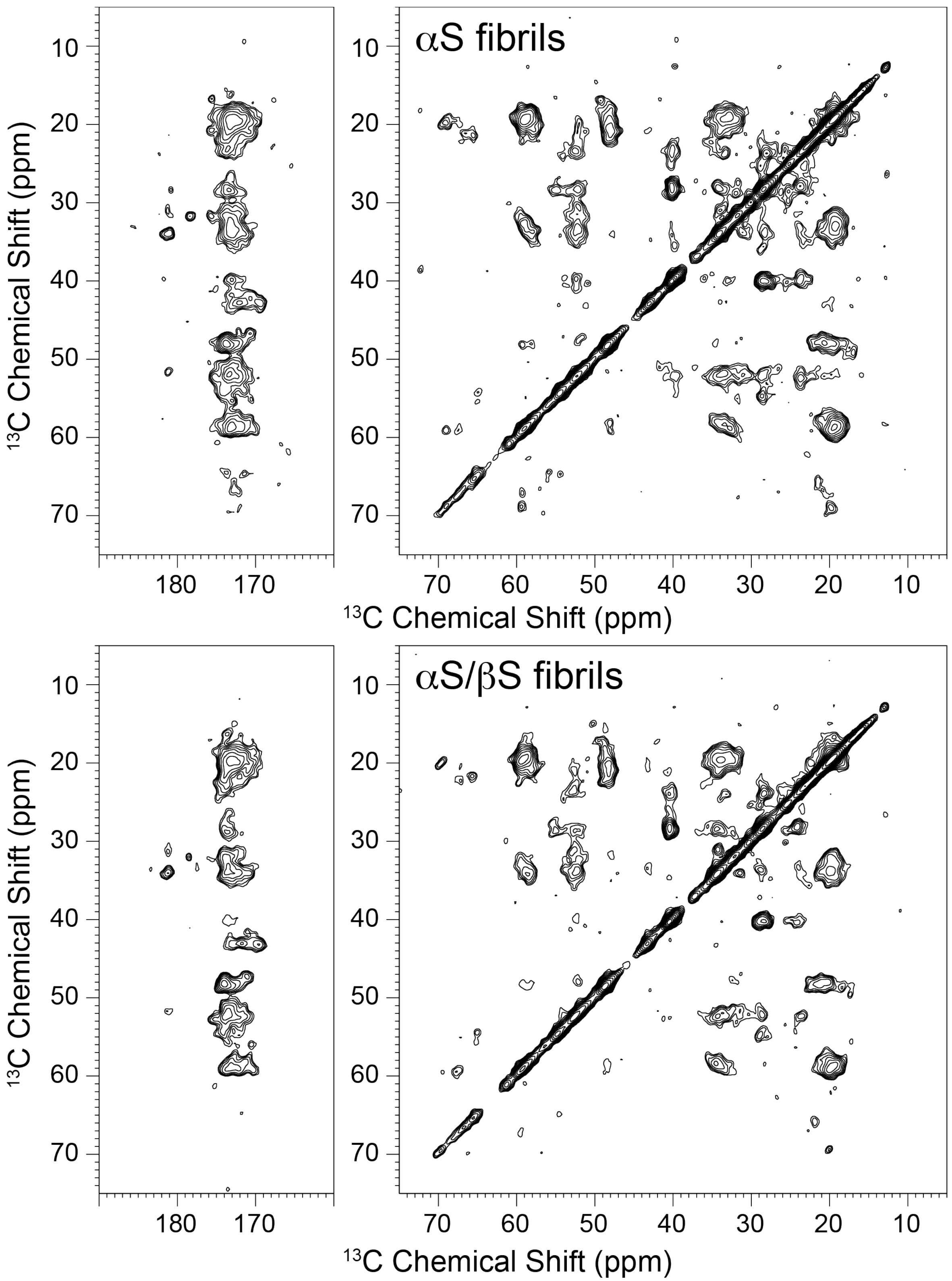


**Figure S4:** Two-dimensional (2D) $^{13}C$-$^{13}C$ dipolar-assisted rotational-resonance (DARR) spectra of αS fibrils (top) and co-incubated αS/βS fibrils (bottom). Spectra were recorded with a DARR mixing time of 100 ms, at a MAS rate of 13.333 kHz, and temperature was controlled at 25 °C.

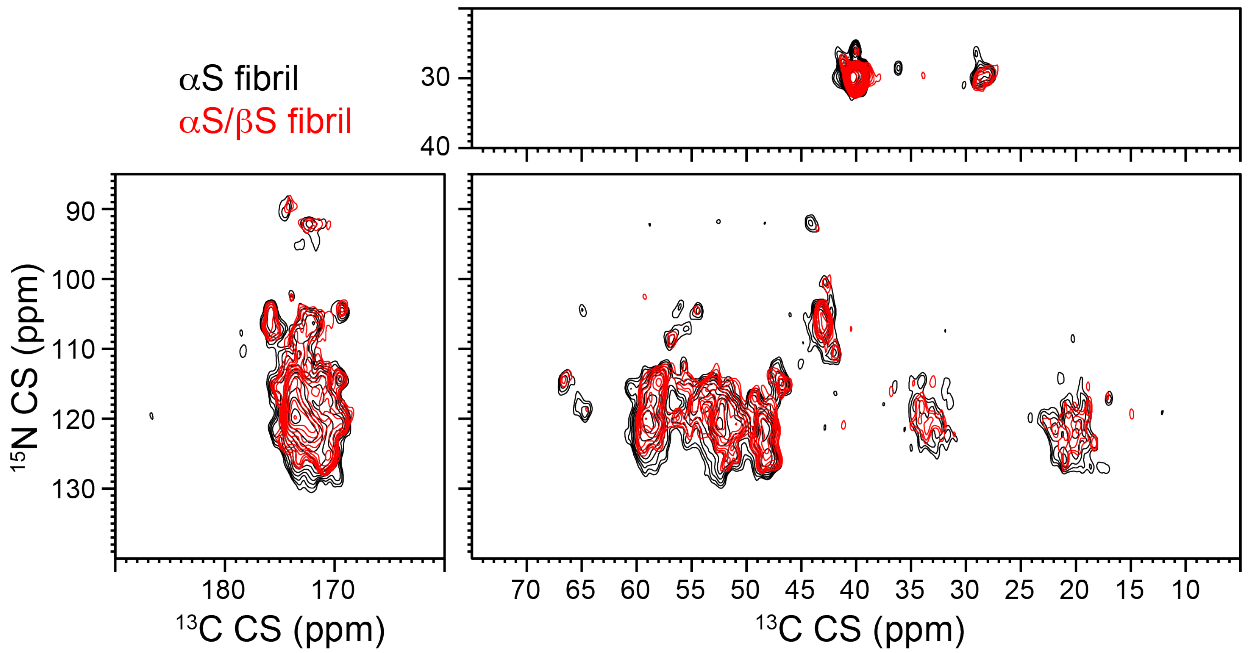


**Figure S5:** Two-dimensional (2D) $^{15}$N-$^{13}$C heteronuclear correlation spectra of αS fibrils (black) and co-incubated αS/βS fibrils (red). Spectra were recorded using a REDOR based pulse sequence with a REDOR period of 1.35 ms, at a MAS rate of 13.333 kHz, and temperature was controlled at 25 °C.

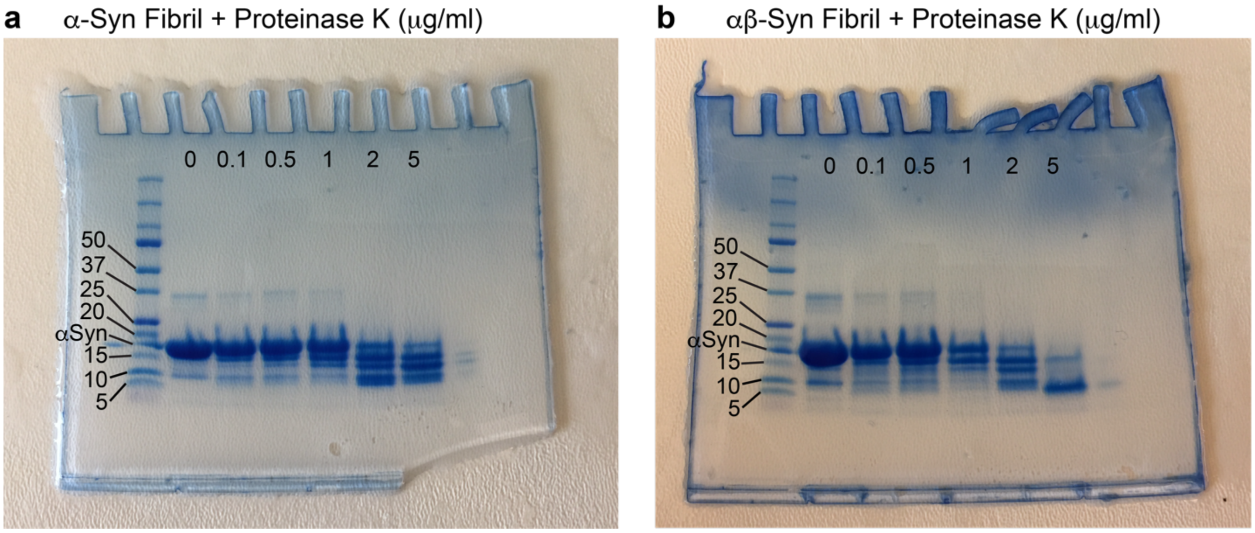


**Figure S6.** Full-length, un-cropped gel images from main text Figure 2c. (a) Digestion of αS fibrils at various concentrations of proteinase K. (b) Digestion of αS/βS fibrils at various concentrations of proteinase K.
